## Supplementary data for "A novel *in-vitro* model of the bone marrow microenvironment in AML identifies CD44 and Focal Adhesion Kinase as therapeutic targets to reverse cell adhesion-mediated drug resistance"

### Supplementary methods

#### Co-culture

The co-culture assay was optimised using different size plates maintaining a similar AML cell to stroma cell ratio and cell density. AML to total stroma was 1.0 $x10^6$ to 1.8 $x10^5$, 5.0$x10^5$ to 9.0$x10^4$, and 8.4$x10^4$ to 1.68 $x10^4$ for 12-, 24- and 96-well plates respectively. On day 1, the plate was coated with 0.2% gelatin for 30 mins. The stroma cell lines cells were plated in DMEM + 10% FBS + 1% P/S + 1% L-glutamine comprising of 1/3 of each of HS5, hFOB 1.19 and HUVEC and left in the incubator overnight to allow confluency. On day 2, the supernatant of each well was removed, and AML cells were added in fresh RPMI + 20% FBS + 1% P/S + 1% L-glutamine ±drugs. For each drug, a dose response curve and optimal pre-incubation time with AML cells was pre-established (supplementary table 2). When cytarabine was used (Selleckchem), it was added after AML cells were cultured in BMAS for 3 hours.

#### Cell adhesion assays

#### BMAS

Non-adhered AML cells in co-culture supernatant were counted, therefore adhesion was assessed indirectly. This was done to negate the need for trypsinization and AML cell selection prior to counting. When using a 24-well plate, cells in the supernatant were counted on day 2, using a Countess II instrument, following vigorous pipetting with a 1mL pipette and trypan blue staining. Each condition was performed in triplicate and the mean number of non-adhered cells calculated. When using the 96-well plate, on day 2 the whole plate was inserted into the CytoFLEX LX where it was agitated for 5 secs before the cell count in 50µL of the supernatant of each well was quantified. The total cell count was then calculated for a 200µL volume. Each condition was performed in triplicate and the mean number of non-adherent cells calculated. Cell lines and primary cells were used in this assay.

**Autologous primary BMAS**

Mononuclear cells were harvested from BM aspirates of AML patients by density gradient centrifugation through Lymphoprep (Stem Cell Technologies, Vancouver, Canada) and cryopreserved in 90% FCS/10% DMSO until required. Layers of autologous stromal cells were generated from the BM aspirates of 3 AML patients using the first step in a previously described methodology (1). One vial of BM mononuclear cells from each patient was thawed and cultured in alpha-MEM containing 10% foetal calf serum, 10mM HEPES, 2U/ml heparin, 250 μM sodium ascorbate, 1mM sodium pyruvate and 2mM Glutamax, in T75 flasks until confluent. After approximately 14 days, the ~80% confluent stromal layers were harvested using Tryple-Express (Thermo Fisher Scientific, Waltham, MA, USA) and 50,000 cells seeded per well of a 96 well plate. After an overnight culture, confluence of the stromal cells was confirmed by microscopy. A second vial of cells from each patient was thawed and $2.5 \times 10^5$ BM mononuclear cells were seeded per well of a 96 well plate in the absence of stromal cells in DMEM media containing 10% FCS, 1g/L D-glucose and 2mM L-glutamine. The cells were cultured for 18 h, either without drug, with anti-CD44, defactinib or combinations of both drugs. The BM mononuclear cells were gently harvested from each well and added to the wells containing the confluent stromal cell layers + and – drugs following aspiration of the media. Data acquisition was performed as described for the cell line BMAS model, with the exception that following the 3h co-culture period, the plate was gently agitated for 30 mins to suspend the non-adherent cells, which were then removed and stained with Live/Dead Aqua (Thermo Fisher Scientific) and an anti-CD34 PE-conjugated antibody (Becton

Dickinson). Following aspiration of the non-adherent cells, the plate was visually inspected to confirm integrity of the stromal cell layer. Data acquisition was performed on a CytoFLEX S flow cytometer (Beckman Coulter), with 2 mins of data collected at a fixed flow rate, per sample. Data are expressed as the fold change in the number of total and $CD34^+$ viable cells present relative to untreated control samples for each patient.

**Isolation of adhered cells from BMAS.**

Adhered cells were isolated following removal of the supernatant containing the non-adherent cells using a pipette. The remaining adhered AML cells and stroma mix were washed 3x times with PBS. TryPLE (Thermofisher) was added and the plate incubated at 37$^o$C until the adherent cell layer detached. Once cells detached, 10% PBS containing 10% FBS was added, and cells were harvested by centrifugation. Volumes of TryPLE used were dependent on the size of the plate following the manufacturer instructions. Isolated cells were stained for phenotyping, apoptosis, cell cycle and ROS levels.

**RNA sequencing and analysis**

Following isolation of the AML cells from the stromal cells, they were resuspended in 1mL ice cold PBS and transferred to pre-cooled RNAse-free Eppendorf tubes, spun at 800g for 5mins at 4$^o$C. Supernatants were removed, pellets snap frozen in liquid nitrogen and then shipped to Active Motif, where RNA was isolated using the MagMAX™ mirVana™ Total RNA Isolation Kit. RNA quality was checked by RNA Screen tape and an RNA integrity number (RIN) score of >8.3 was used as a threshold. The samples were all treated with DNase. Library preparation was performed with TruSeq stranded mRNA library kit (Illumina). Analysis was performed using the free online software BioJupies[1].

**RT-PCR validation for RNA sequencing**

RNA quantification

RNA which was isolated from samples by Active Motif (see 3.13) was subsequently shipped back and stored at -80°C. RNA quality and concentration was measured three times using a Nanodrop spectrophotometer ND1000 and the mean concentration was calculated for each technical replicate. A 260nm/280nm absorbance ratio >1.8 was deemed acceptable.

Reverse Transcription

Reverse transcription was performed using High-Capacity RNA-to-cDNA™ Kit. 1μg RNA was added in a 10μL reaction mix (5μL 2X RT buffer mix, 0.5μL 20X RT enzyme mix, up to 4.5μL RNA sample and nuclease-free $H_2O$ to make up to a total of 10μL). Tubes were briefly centrifuged to spin down the contents and to eliminate any air bubbles. Samples were incubated at 37°C for 60 minutes. The reaction was subsequently stopped by heating the samples to 95°C for 5 minutes and then holding them at 4°C. This was carried out using the a GeneAmp PCR System 9700 (Thermo Fisher). The cDNA was stored at −20°C.

Quantitative real-time PCR

Real Time quantitative PCR was performed in 96 well plates in triplicate for each condition. Each reaction contained 5μL of TaqMan™ fast advanced master mix (2X), 0.5μL of TaqMan™ Assay (20X), 3.5μL of nuclease-free water and 1μL of cDNA. The reaction plate was sealed with optical adhesive film, then briefly centrifuged. Real-time

PCRs were run on a ViiA™ 7 Real-Time PCR System (Applied Biosystems) and the resulting data were analysed using the ΔΔCt method. Briefly, the average Ct value of the triplicates was calculated. The ΔCt value was calculated by subtracting the Ct value of the *ACTB* housekeeping gene from the Ct value of the gene of interest (Ct gene of interest-Ct *ACTB*). The ΔΔCt value was calculated by subtracting the ΔCt value of the control (untreated cells) from the ΔCt value of treated cells (ΔCt treated-ΔCt untreated). The fold difference between treated and untreated samples was then calculated as 2-ΔΔCt.

**Measuring pFAK**

The levels of pFAK was were measured using the True-Phos™ kit (Biolegend) and PE-anti-pFAK (BD Biosciences) as per the manufacturers instructions. The MFI values for p-FAK were determined in gated CD45+ CD73- AML cells.

**Statistical analysis and synergy**

All statistical analyses were performed using GraphPad Prism 9.5.1 (GraphPad Software, San Diego, CA, USA). In all cases, the normal distribution of the data was assessed using the Shapiro–Wilk test. Assuming that this assumption was met, univariate comparisons were made using a t-test for paired observations comparing the mean of the technical triplicates between the two groups. If the means between > 2 groups were compared, one-way ANOVA with Dunnett's correction (which compares every mean to a control mean) for multiple comparisons was used. Toxicity data from the drug treatments were used to produce sigmoidal dose–response curves. From these interpolated $LC_{50}$ values (the concentration of drug required to kill 50% of the cells) were calculated following normalisation and constraint (bottom value = 0 and top value between 0 and 100). The expected drug combination responses were calculated based on a ZIP reference model using SynergyFinder[27]. Deviations between observed and expected responses with positive and negative values denote synergy and antagonism respectively. For estimation of outlier measurements, the cNMF algorithm[28] implemented in SynergyFinder was utilized. For all statistical tests a confidence level of 95% was used, and therefore p values of <0.05 were deemed significant.

1. Reinisch A, Hernandez DC, Schallmoser K, Majeti R. Generation and use of a humanized bone-marrow-ossicle niche for hematopoietic xenotransplantation into mice. Nat Protoc. 2017;12(10):2169-88.

| BSMS Patient ID | Sample | Diagnosis | Gender | Age | Risk (ELN) |
|---|---|---|---|---|---|
| 1 | BM/PB | Acute myelomonocytic leukaemia | F | 78 | Adverse |
| 2 | BM/PB | AML without Maturation | F | 28 | Intermediate |
| 3 | BM/PB | AML<br>With CEBPA mutation | M | 71 | Favourable |
| 4 | BM/PB | AML with RUNX1::RUNX1T1 fusion | M | 49 | Intermediate |
| 5 | BM/PB | AML with multilinear dysplasia (MLD) | M | 41 | Adverse |
| 6 | BM/PB | AML (previously B BPDCN) | M | 64 | Adverse |
| 7 | BM/PB | AML | F | 21 | N/A |
| 8 | BM/PB | AML with mutated NPM1 (AML with recurrent genetic abnormalities) | F | 74 | Intermediate |
| 9 | BM | Acute Myeloid Leukaemia with Mutated RUNX1 | M | 65 | Adverse |
| 10 | PB | Secondary AML (Previous CMML) | M | 82 | Adverse |
| 11 | PB | Secondary AML (previous ET) | F | 61 | Adverse |
| 12 | PB | AML with KMT2A (MLL) (11q23) rearrangement | M | 63 | Adverse |
| 13 | PB | Acute myeloid leukaemia, myelodysplasia-related | F | 60 | Adverse |
| 14 | PB | Acute myeloid leukaemia, myelodysplasia-related | F | 77 | Adverse |
| 15 | PB | No bone marrow done, analysis on PB only | M | 65 | Adverse |
| **Flinders Patient ID** | | | | | |
| AML13 | BM | AML with<br>47,XY,+11[9]/46,XY[1]<br>FLT3-ITD positive<br>IDH2 p.(Arg140Gln) | M | 78 | Adverse |
| AML15 | BM | AML with normal karyotype<br>NPM1 p.(Trp288CysfsTer12) | M | 55 | Favourable |
| AML17 | BM | AML sample after induction failure<br>Normal karyotype<br>IDH2 p.(Arg140Gln) | F | 61 | Intermediate |

**Supplementary Table 1: Clinical characteristics of newly diagnosed patients, who donated PB and/or BM samples.**

| Release agent | Target | Supplier | Optimal pre-incubation time |
|---|---|---|---|
| Plerixafor | CXCR4 | Generon | 1 hour |
| ONO7161 | CXCR4 | ONO Pharmaceutical Ltd. | 1 hour |
| Tysabri (Natalizumab) | alpha-4 integrin (CD49d) | Biogen Inc. | 1 hour |
| Purified NA/LE Mouse Anti-Human CD44 (Clone 515) | CD44 | BD Pharmingen™ | 1 hour or 18 hours (in combination experiments) |
| Anti-human E-selectin/CD62E Antibody | E-selectin | R&D Systems | 1 hour |
| Defactinib hydrochloride | FAK and PYK2 | Bioscience | 18 hours |

**Supplementary Table 2: Release agents tested.**

| | | OCI-AML3 | | KG1a | | HS-5 | | hFOB 1.19s | | HUVEC | |
|---|---|---|---|---|---|---|---|---|---|---|---|
| | **Supplier and clone** | **%** | **MFI** | **%** | **MFI** | **%** | **MFI** | **%** | **MFI** | **%** | **MFI** |
| **CD45** | Biolegend H130 | 100 | 262998 | 100 | 283674 | 1 | -482 | 3 | -571 | 58 | 7478 |
| **CD34** | Biolegend WM53 | 85 | 30599 | 100 | 71062 | 2 | 9384 | 3 | 7855 | 2 | 10762 |
| **CD73** | Biolegend AD2 | 98 | 11600 | 0 | 638 | 100 | 69481 | 98 | 53246 | 100 | 82077 |
| **CD44** | Biolegend BJ18 | 100 | 80141 | 100 | 102948 | 100 | 265978 | 100 | 284238 | 100 | 392896 |
| **CXCR4** | Biolegend 12G5 | 100 | 181099 | 100 | 53596 | 0 | -1117 | 6 | 1405 | 100 | 28717 |
| **CD49d** | Biolegend 9F10 | 100 | 25525 | 100 | 28834 | 4 | 19398 | 25 | 167112 | 37 | 25225 |
| **E selectin** | Biolegend HAE-1f | 2 | 1035 | 5 | 869 | 1 | 3071 | 1 | 2600 | 21 | 6607 |

**Supplementary Table 3: Percentage (%) of cells with expression and MFI of different surface antigens on the two AML cell lines, HS-5, hFOB 1.19 and HUVEC.**

| | | |
|---|---|---|
| AC 0.94 vs. B 0.94 | **** | <0.0001 |
| AC 0.94 vs. B 1.25 | **** | <0.0001 |
| AC 0.94 vs. B 2.50 | **** | <0.0001 |
| AC 0.94 vs. B 5 | *** | 0.0002 |
| AC 1.25 vs. B 0.94 | **** | <0.0001 |
| AC 1.25 vs. B 1.25 | **** | <0.0001 |
| AC 1.25 vs. B 2.50 | **** | <0.0001 |
| AC 1.25 vs. B 5 | *** | 0.0002 |
| AC 2.50 vs. B 0.94 | **** | <0.0001 |
| AC 2.50 vs. B 1.25 | **** | <0.0001 |
| AC 2.50 vs. B 2.50 | **** | <0.0001 |
| AC 2.50 vs. B 5 | **** | <0.0001 |
| AC 5 vs. B 0.94 | **** | <0.0001 |
| AC 5 vs. B 1.25 | **** | <0.0001 |
| AC 5 vs. B 2.50 | **** | <0.0001 |
| AC 5 vs. B 5 | **** | <0.0001 |
| D 0.94 vs. B 0.94 | **** | <0.0001 |
| D 0.94 vs. B 1.25 | **** | <0.0001 |
| D 0.94 vs. B 2.50 | **** | <0.0001 |
| D 0.94 vs. B 5 | **** | <0.0001 |
| D 1.25 vs. B 0.94 | **** | <0.0001 |
| D 1.25 vs. B 1.25 | **** | <0.0001 |
| D 1.25 vs. B 2.50 | **** | <0.0001 |
| D 1.25 vs. B 5 | **** | <0.0001 |
| D 2.50 vs. B 0.94 | **** | <0.0001 |
| D 2.50 vs. B 1.25 | **** | <0.0001 |
| D 2.50 vs. B 2.50 | **** | <0.0001 |
| D 2.50 vs. B 5 | **** | <0.0001 |
| D 5 vs. B 0.94 | **** | <0.0001 |
| D 5 vs. B 1.25 | **** | <0.0001 |
| D 5 vs. B 2.50 | **** | <0.0001 |
| D 5 vs. B 5 | **** | <0.0001 |

**Supplementary Table 4:** Significance values for different concentrations of anti-CD44 alone (AC) and defactinib alone (D) versus the combination of both (B). Concentrations; 0.94µg/mL, 1.25µg/mL, 2.5µg/mL and 5µg/mL. Significance determined by one-way ANOVA and Dunnett's multiple comparisons test, following Shapiro-Wilk test for normality, ****p ≤ 0.0001, ***p ≤ 0.001.

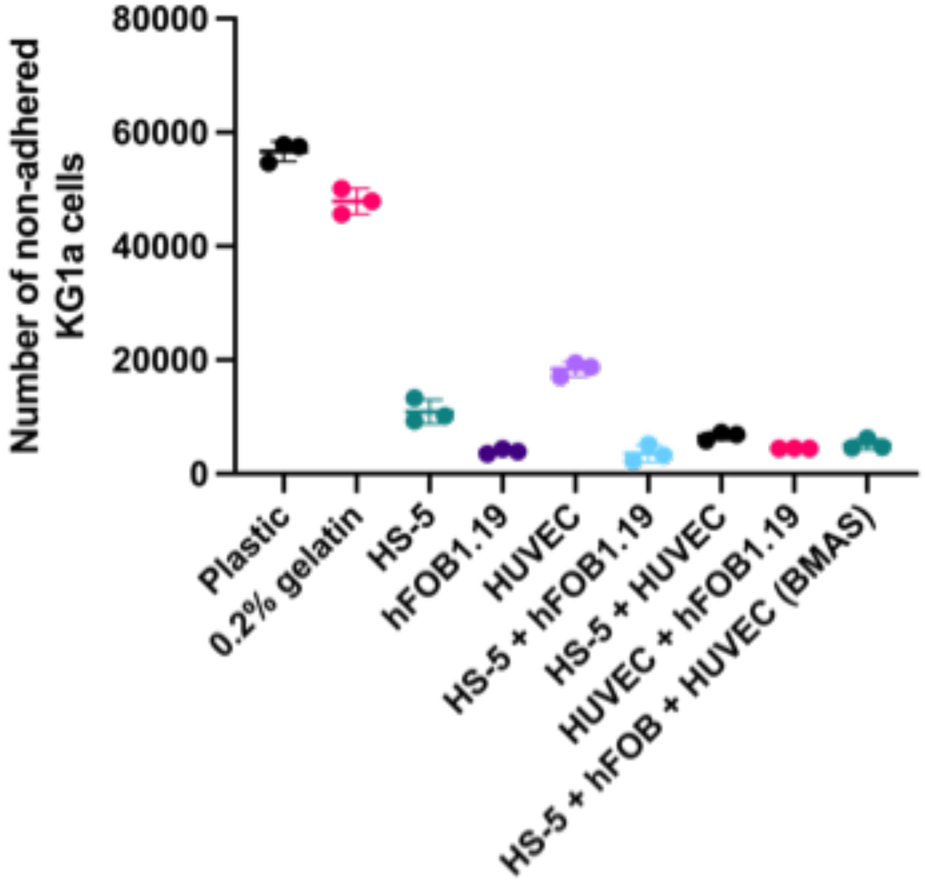


**Supplementary Figure 1: Adhesion of KG1a cells to different stromal cell combinations indicates that hFOB 1.19 cells form the most adhesive stromal layer.** KG1a cells (84,000 cells per well) were added to wells pre-seeded 24 hours earlier with 18,600 stromal cells of different types (HS-5, hFOB1.19, or HUVEC) or their combinations in equal ratios. The number of non-adhered KG1a cells was quantified 3 hours post-incubation as an indirect measure of the adhesive properties of each condition. All wells, except those in the plastic-only control, were coated with 0.2% gelatin as it was used to facilitate HUVEC adhesion to plastic. This gelatin coating had minimal impact on KG1a cell adhesion, making the 0.2% gelatin condition a baseline for assessing the adhesive capacity of the stromal cells themselves. Data presented as mean ± SD of 3 technical replicates.

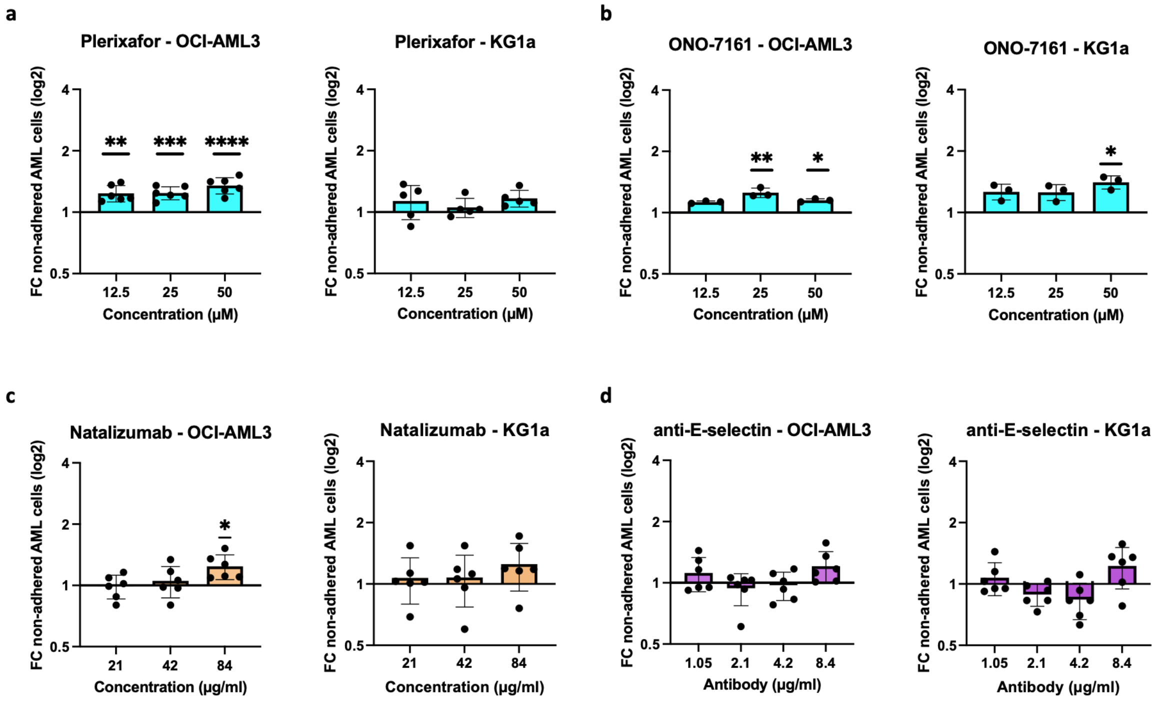


**Supplementary Figure 2: Incubation of AML cells with plerixafor, ONO-7161, natalizumab and anti-E-selectin on the BMAS.** (a) Fold change (FC) in non-adhered AML cells (mean±SD) when treated with increasing doses of plerixafor and then co-cultured for 3 hours in OCI-AML3 (n=6) and KG1a (n=6). (b) Fold change (FC) in non-adhered AML cells (mean±SD) when treated with increasing doses of ONO-7161 and then co-cultured for 3 hours in OCI-AML3 (n=6) and KG1a (n=6) cells. (c) Fold change (FC) in non-adhered OCI-AML3 (n=6) and KG1a (n=6) AML cells (mean±SD) when treated with increasing doses of natalizumab and then co-cultured for 3 hours. (d) Fold change (FC) in non-adhered OCI-AML3 (n=6) and KG1a (n=6) AML cells (mean±SD) when treated with increasing doses of E-selectin antibody and then co-cultured for 3 hours. Significance determined by one-way ANOVA and Dunnett's multiple comparisons test (comparing every mean to a no treatment control which is equal to 1), following Shapiro-Wilk test for normality, ****$p \leq 0.0001$, ***$p \leq 0.001$, **$p \leq 0.01$, *$p \leq 0.05$.

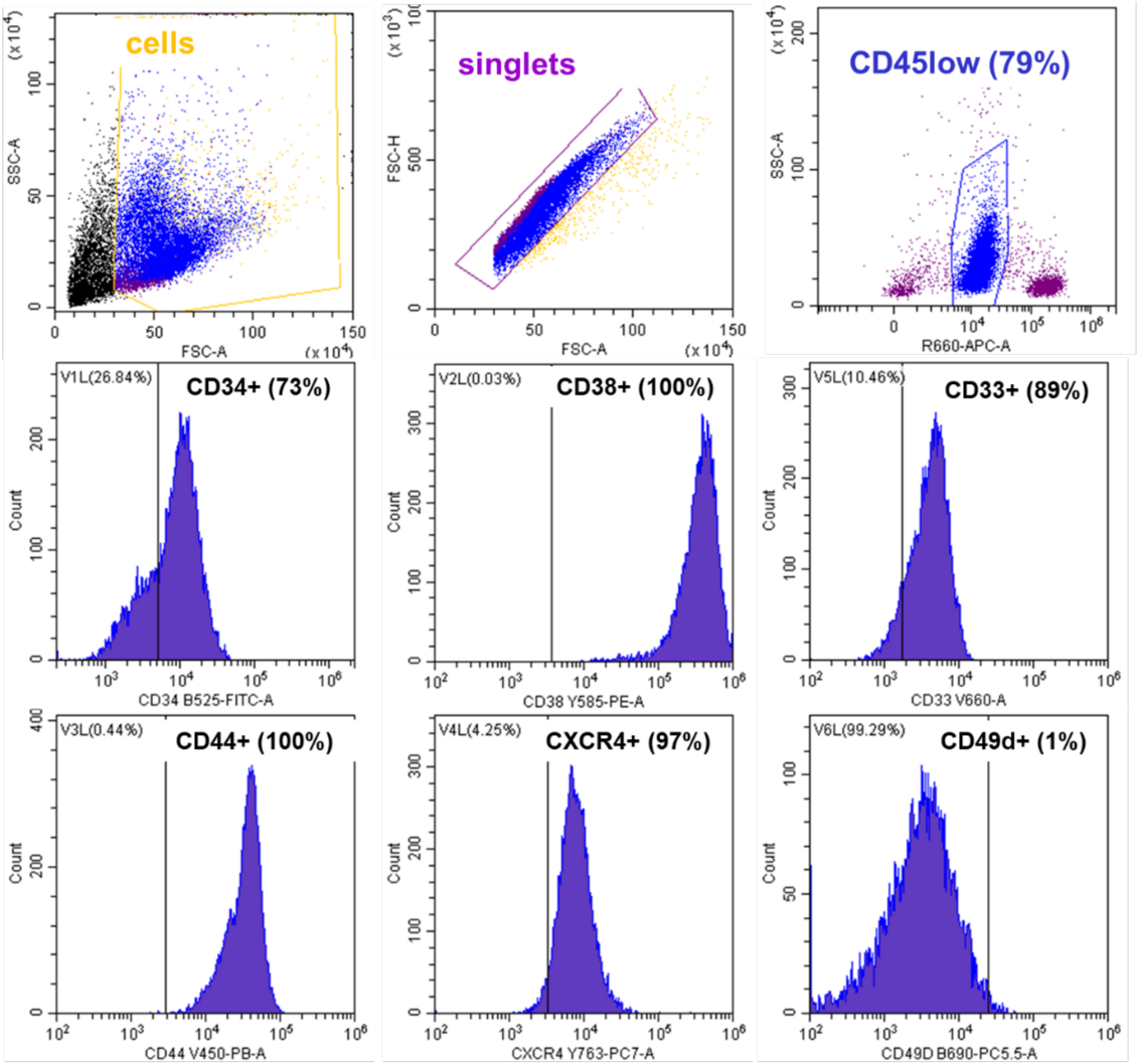


**Supplementary Figure 3: Gating strategy for primary AML cells.** Representative dot plots and histograms from one AML patient sample illustrating the gating strategy to identify blasts as CD45$^{low}$ using a CD45/SSC plot. CD45$^{-}$ (nucleated erythrocytes) and CD45$^{high}$ (lymphocytes) cells were excluded from analysis. Expression of the other cell surface markers was assessed in the CD45$^{low}$ blast population (blue).

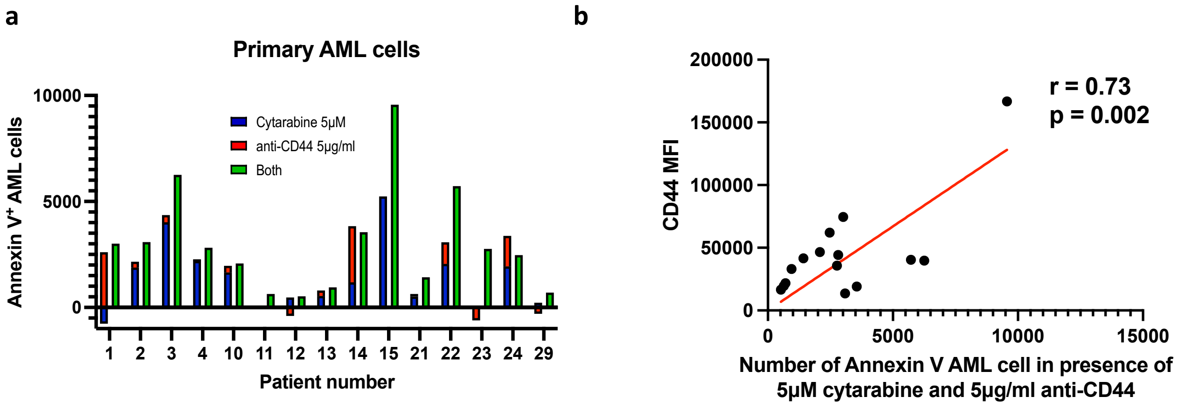


**Supplementary Figure 4: The combination of anti-CD44 with cytarabine can overcome CAM-DR better than the the summative effect of either alone.** (a) Number of Annexin V positive AML cells (in 50ul) following treatment with 5µM of cytarabine alone (blue), 5µg/mL anti-CD44 alone (red) in primary AML cells. The sum of their individual effect (red/blue column) is compared to their combination effect when cells were treated with both agents simultaneously (green column). (b) The number of Annexin V positive AML cells in the presence of anti-CD44 and cytarabine correlates with their pre-treatment expression of CD44 As determined by Pearson's correlation.

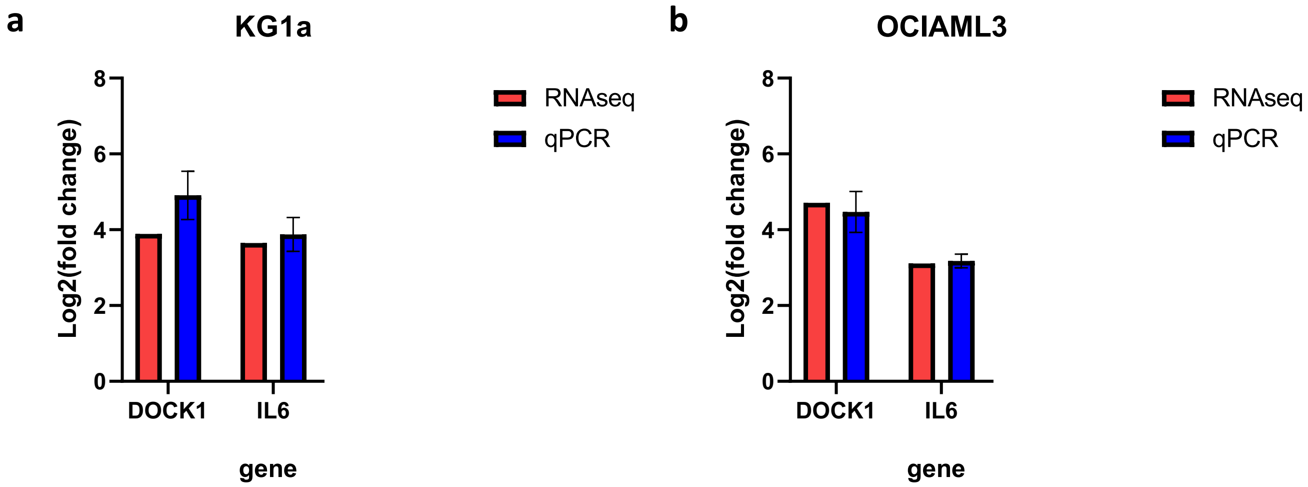


**Supplementary Figure 5: Validation of RNA sequencing results were validated using RT-PCR.** Two genes that were part of the FAK signalling pathway were chosen, namely *DOCK1* and *IL6*. RNA sequencing data (red) from KG1a (n=3) (a) and OCI-AML3 (n=3) cells (b). qPCR data (blue) (n=3 per cell line) as assessed by a TaqMan assay.

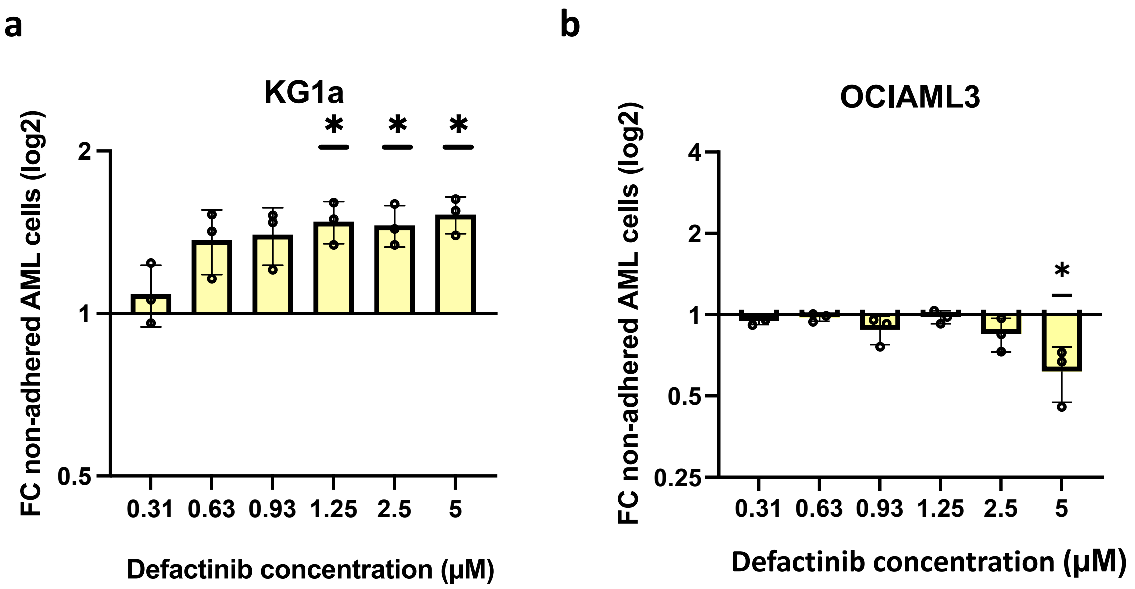


**Supplementary Figure 6: Defactinib blocked adhesion in the more primitive KG1a cells did not in the more differentiated OCI-AML3 cells.** (a) Fold change (FC) in non-adhered KG1a cells (mean±SD) following treatment with increasing doses of defactinib and a 3 hour co-culture (n=3). (b) Fold change (FC) in non-adhered OCI-AML3 cells (mean±SD) following treatment with increasing doses of defactinib and a 3 hour co-culture (n=3). Significance determined by one-way ANOVA and Dunnett's multiple comparisons test (comparing every mean to a no treatment control which is equal to 1), following Shapiro-Wilk test for normality, *$p \leq 0.05$.

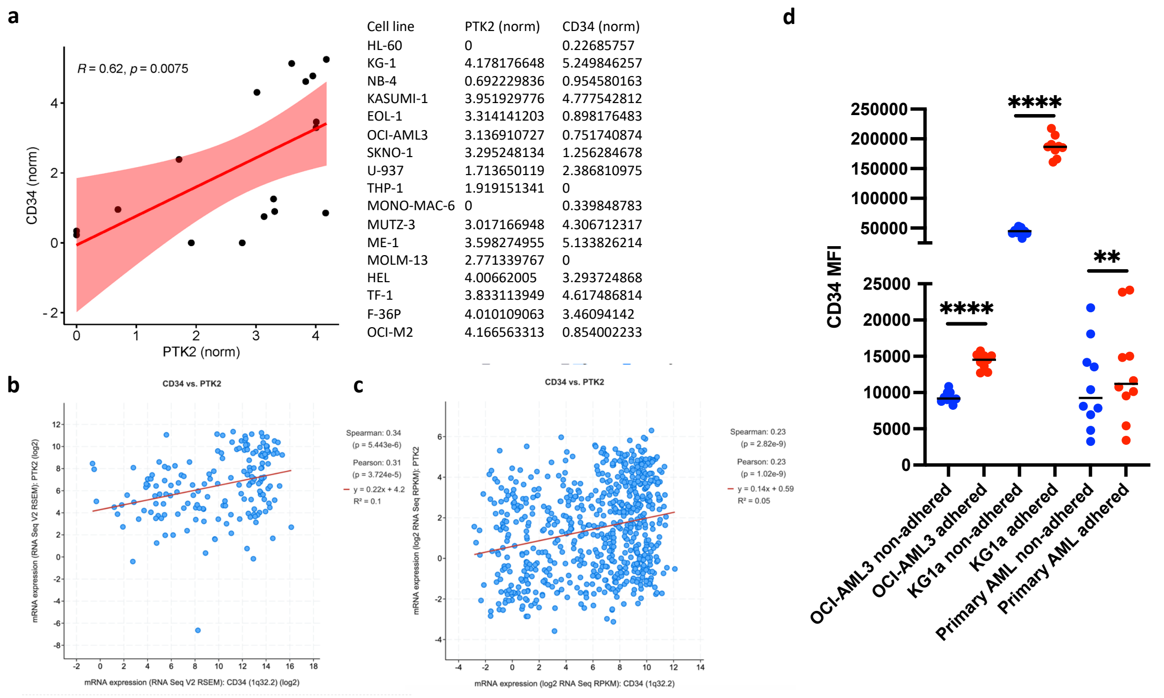


| Cell line | PTK2 (norm) | CD34 (norm) |
|---|---|---|
| HL-60 | 0 | 0.22685757 |
| KG-1 | 4.178176648 | 5.249846257 |
| NB-4 | 0.692229836 | 0.954580163 |
| KASUMI-1 | 3.951929776 | 4.777542812 |
| EOL-1 | 3.314141203 | 0.898176483 |
| OCI-AML3 | 3.136910727 | 0.751740874 |
| SKNO-1 | 3.295248134 | 1.256284678 |
| U-937 | 1.713650119 | 2.386810975 |
| THP-1 | 1.919151341 | 0 |
| MONO-MAC-6 | 0 | 0.339848783 |
| MUTZ-3 | 3.017166948 | 4.306712317 |
| ME-1 | 3.598274955 | 5.133826214 |
| MOLM-13 | 2.771339767 | 0 |
| HEL | 4.00662005 | 3.293724868 |
| TF-1 | 3.833113949 | 4.617486814 |
| F-36P | 4.010109063 | 3.46094142 |
| OCI-M2 | 4.166563313 | 0.854002233 |

**Supplementary Figure 7: Expression of CD34 correlates with FAK (PTK2) and increased adhesion in AML.** (a) Analyses of previously published CD34 versus FAK (PTK2) mRNA levels in cell lines classified as AML myelocytic/monocytic/erythroid from the LL-100 cell line dataset (Quantmeier et al: doi.org/10.1038/s41598-019-44491-x). A pseudocount of 1 was added to normalised counts prior to log2 transformation. (b and c) Analysis of previously published CD34 versus FAK (PTK2) mRNA levels in two AML patient cohorts (b, Ley et al: doi: 10.1056/NEJMoa1301689 and c, Bottomly et al: doi.org/10.1016/j.ccell.2022.07.002.). Patient data was analysed using cBioPortal (Cerami et al: DOI: 10.1158/2159-8290.CD-12-0095.) (d) Mean Fluorescence Intensity (MFI) of CD34 expression between adhered versus non-adhered OCI-AML3 (n=9), KG1a cells (n=9) and primary AML cells (n=10). Significance was determined using a paired t-test, following Shapiro-Wilk test for normality. **** $p < 0.0001$, *** $p < 0.001$** $p < 0.01$.
